# Extensive and deep sequencing of the Venter/HuRef genome for developing and benchmarking genome analysis tools

**DOI:** 10.1101/281709

**Authors:** Bo Zhou, Joseph G. Arthur, Steve S. Ho, Reenal Pattni, Yiling Huang, Wing H. Wong, Alexander E. Urban

## Abstract

We produced an extensive collection of deep re-sequencing datasets for the Venter/HuRef genome using the Illumina massively-parallel DNA sequencing platform. The original Venter genome sequence is a very-high quality phased assembly based on Sanger sequencing. Therefore, researchers developing novel computational tools for the analysis of human genome sequence variation for the dominant Illumina sequencing technology can test and hone their algorithms by making variant calls from these Venter/HuRef datasets and then immediately confirm the detected variants in the Sanger assembly, freeing them of the need for further experimental validation. This process also applies to implementing and benchmarking existing genome analysis pipelines. We prepared and sequenced 200 bp and 350 bp short-insert whole-genome sequencing libraries (sequenced to 100x and 40x genomic coverages respectively) as well as 2 kb, 5 kb, and 12 kb mate-pair libraries (49x, 122x, and 145x physical coverages respectively). Lastly, we produced a linked-read library (128x physical coverage) from which we also performed haplotype phasing.

## BACKGROUND & SUMMARY

Almost two decades ago the extensive efforts of the Human Genome Project, backed up by work from Celera, resulted in the release of a draft of the first complete sequence of the human genome ^1,2^. This catalyzed a new era of human whole-genome analysis where the now-available human genome sequence has been studied intensely to understand the functions of its parts and their interactions with each other and where a concurrent genome technology revolution has produced ever more powerful platforms to carry out such functional studies ^3^. Since then, increasingly large numbers of human genomes have been sequenced, yielding insights into population-level genetic variation ^4–6^, structural genome variation ^7–9^, and mutational mechanisms ^10^. Technological advances have progressively improved the information content and reduced the noise profile of sequencing data ^11^. A large variety of methodologies for the routine analysis of sequencing data is now available ^12^. “Whole-genome sequencing” is now a standing term that refers to the re-sequencing of a given sample of human genomic DNA using, typically, the dominant Illumina DNA sequencing platforms which can quickly produce several hundred million short sequencing reads at affordable costs. These reads are then aligned to the human reference genome and analyzed using various approaches ^12–14^, such as mismatch analysis, read-depth analysis, split-read analysis and discordant read-pairs analysis, producing an extensive catalog of sequence variants that are present in the DNA sample in question relative to the human reference sequence. The promise of human genome research is nothing short of a complete transformation of basic life science research, translational research, and eventually the way we diagnose, treat, and find cures for human disease.

It is clear, however, that current standard whole-genome sequence analysis leaves a rather large room for improvement. The standard genome analysis practices of today perform rather poorly in certain contexts, such as in repetitive regions (i.e. in around half the human genome), in the detection and resolution of complex structural variation, or in placing detected variants in their proper haplotypes. Although more advanced and novel computational algorithms that address these limitations are continuously being developed, one essential requirement during this process is that the detected variants are to be experimentally validated in order to establish false-positive rates and to make it possible to further tune and optimize the new algorithms. Experimental validation, especially of complex variants, during the tool development and testing phases is a very laborious and time-consuming process, but it can be circumvented by using a genome for which sufficiently large numbers of variants are already known, i.e. prevalidated. Several studies have been conducted with the goal of extensively characterizing the variants in a small number of human genomes using multiple sequencing technologies ^15,16^. In some human genomes, variants have been carefully and extensively documented, providing a benchmark for other studies ^9,17–20^.

The Venter (HuRef) Genome, however, is especially distinguished for quality among the publicly-available human genome sequences as it is the only one for which its complete diploid assembly was generated from high-quality Sanger reads ^17^ and for which extensive catalogs of SNPs, indels, and structural variation are available ^18,20^. To date, no extensive Illumina sequencing datasets have been available for the Venter/HuRef genome in contrast to other genomes that have been characterized for benchmarking purposes ^15,16^.

To unlock the potential of the Venter/HuRef genome as the outstanding benchmark genome, we have conducted deep whole-genome sequencing (WGS) using a variety of sequencing strategies for the Illumina platform (**Table 1**). Specifically, we produced short-insert paired-end WGS datasets at a combined sequence coverage of 140x, linked-read data at 42x de-duplicated sequencing coverage (128x physical coverage i.e. the average number of times the genome is spanned by input DNA fragments rather than the average number of times the genome is covered by sequencing reads as in sequencing coverage), and three long-insert (2 kb, 5 kb, and 12 kb) paired-end (i.e. mate-pair) WGS datasets with physical coverages of 49x, 122x, and 145x, respectively (**Figure 1**). These datasets are of very high quality (**Figures 2-4**) and are complemented by the existing Venter/HuRef assembly-quality Sanger reads ^17^ and long-read sequencing data, which was produced using the Pacific Biosciences platform ^21^.

**Figure 1.**
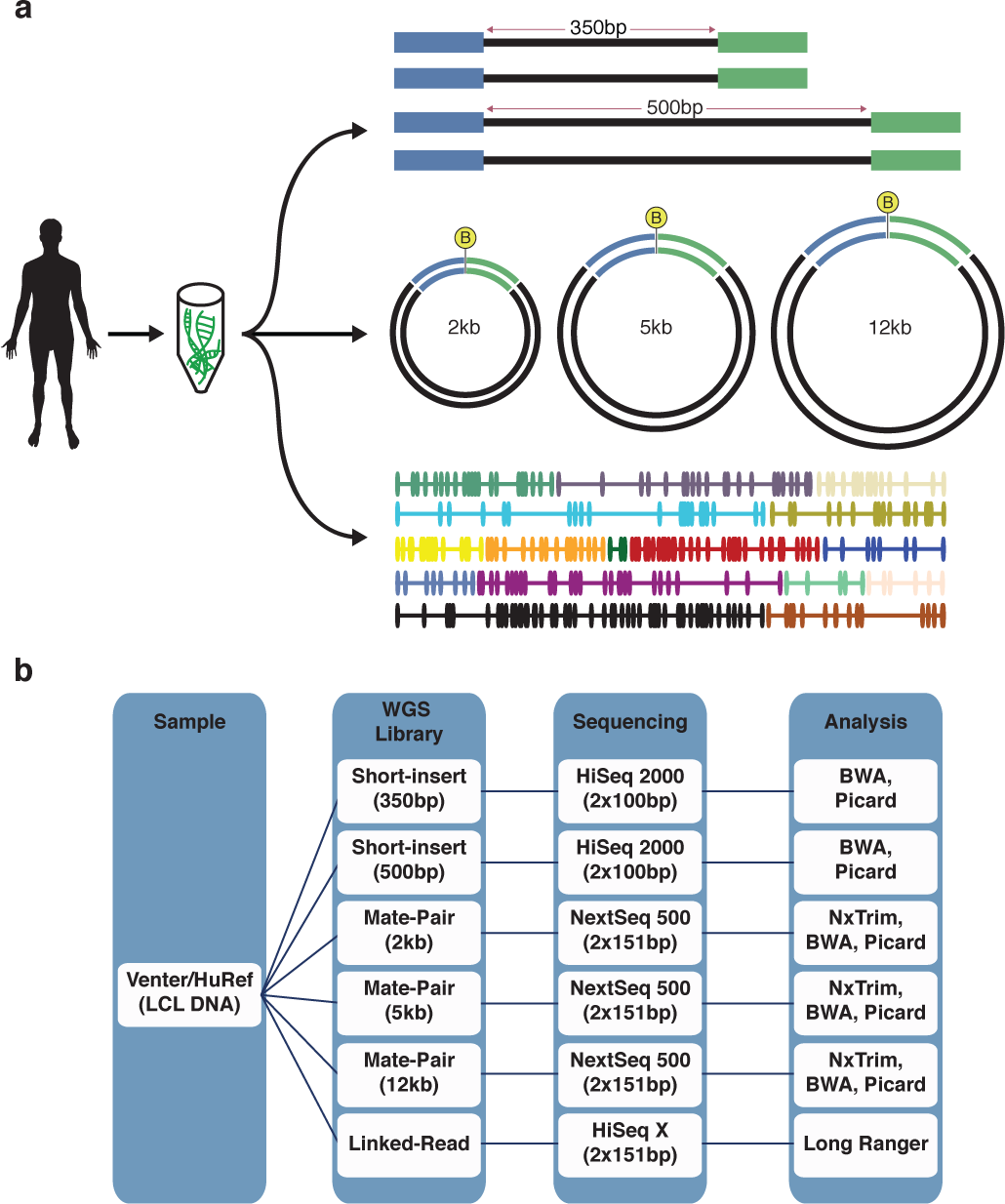
Schematic diagram of the study. **(a)** Venter/HuRef genomic DNA was used to generate short-insert (200 bp and 350 bp), mate-pair (2 kb, 5 kb, and 12 kb), and linked-read libraries. (**b**) Detailed overview of data generation including bio-sample used, types of Illumina WGS libraries constructed, sequencing instrument platforms, types of sequencing runs, and subsequent analysis of data.

**Figure 2.**
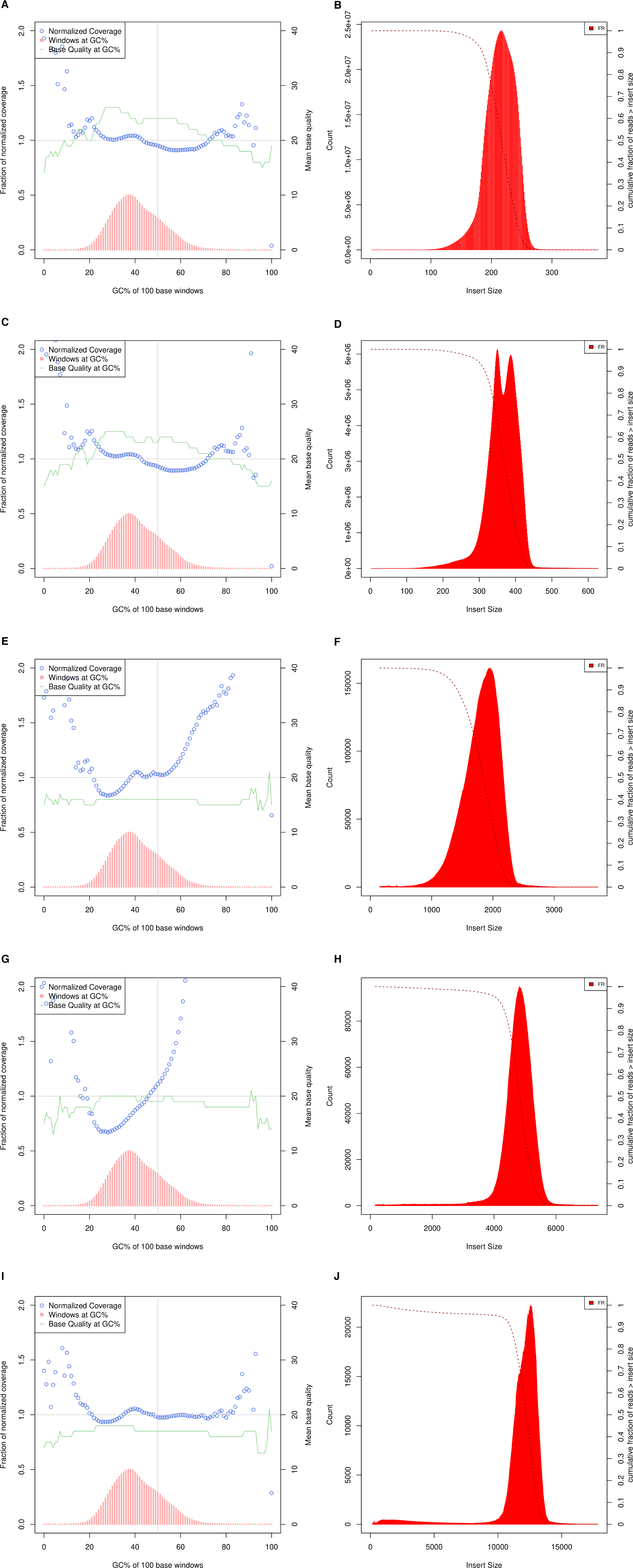
Normalized coverage, GC (%) content windows, base quality at GC (%), and corresponding insert-size histograms for all WGS libraries. **(a, b)** 200 bp short-insert, **(c, d)** 350 bp short-insert, **(e, f)** 2kb-mate-pair, **(g, h)** 5kb-mate-pair, **(i, j)** 12kb-mate-pair.

Researchers developing novel computational tools for analyzing whole-genome sequencing data can now test their algorithms by processing the appropriate Venter/HuRef Illumina datasets described here and then turn to the already-available catalogs of sequence variants, or to the original Sanger reads ^17^, to confirm the characterization of variants detected by their algorithms. Likewise, whenever a laboratory implements a new computational pipeline for human genome analysis, it can now use these Illumina Venter/HuRef datasets to confirm proper implementation and to optimize proper settings for the pipeline.

## METHODS

### Venter/HuRef DNA Sample

The Venter/HuRef DNA sample as obtained as a 50 µg aliquot of LCL-extracted DNA (NS12911) from the Coriell Institute for Medical Research where the iPSC (GM25430) of the same subject is also available (https://catalog.coriell.org/1/HuRef).

### Illumina paired-end WGS

#### Library Preparation

The library preparation was previously described in detail in Mu *et al* ^20^. Briefly, 1 μg of genomic DNA was fragmented using 2μL of NEBNext dsDNA fragmentase (New England Biolabs, Ipswich, MA) in 1x fragmentation buffer and 1x BSA. Reaction was kept on ice for 5 minutes before adding the fragmentase and was incubated at 37□°C for 20 minutes. The reaction was stopped by addition of 5 μL of 0.5 M EDTA. DNA was purified from the reaction mixture using 0.9x by volume AMPure XP beads (Beckman Coulter, Cat# A63880) and eluted in 50 μL of 10mM Tris-Acetate (pH 8.0) buffer. Six independent fragmentation reaction replicates were performed, and the sizes of the DNA were analyzed using Agilent 2100 Bioanalyzer before library preparation.

Library preparation was performed using the KAPA Library Preparation kit (KAPA Biosystems, Wilington, MA) where 200 ng of fragmented DNA was used as input. Library was constructed according to manufacture’s protocol where the DNA was end-repaired and A-tailed before adapter ligation with Illumina TruSeq Adapter (Index 1). DNA was then purified using 0.8x by volume AMPure XP beads and quantified using the Qubit ds DNA High Sensitivity Assay Kit (Life Technologies, Cat# Q32851). For PCR amplification, 50 ng of DNA was amplified using the KAPA HiFi DNA Polymerase with the following thermocycling conditions: 98□°C/45□s, 5 cycles of (98□°C/15□s, 60□°C/30□s, 72□°C/45□s), 72□°C/1□min, and 4□°C /hold. Primers from the KAPA Library Preparation kit was used for PCR amplification. Afterwards, DNA was purified from the PCR reaction using AMPure XP beads and eluted in 30 μL of 10mM Tris-Acetate (pH 8.0) buffer. Six independent experimental replicates were performed, and the purified PCR amplified DNA fragments from each replicate was pooled for size selection and gel-purified from 2% agarose gel. Two size selections were made at 200 bp and 350 bp.

#### Sequencing

Sequencing of the 200 bp and 350 bp insert-size libraries was described previously in Mu et al ^20^. The libraries were sequenced separately (2×100 bp) on an Illumina HiSeq 2000 instrument in rapid run mode. For the 200 bp insert-size library, a total of 3,214,626,588 reads generated from 5 sequencing runs was pooled together to obtain 100x genomic coverage. For the 350 bp insert-size library, a total of 1,280,576,580 reads generated from two sequencing runs was pooled together to obtain 40x genomic coverage.

#### Analysis

Reads were trimmed at the 3’ end to a uniform length of 100 bp using FASTX toolkit (http://hannonlab.cshl.edu/fastx_toolkit/; version 0.0.13). The trimmed reads were aligned by BWA-MEM (Li and Durbin 2009; version 0.7.17-r1188) using the hg38 reference with ALT alleles removed (ftp://ftp.ncbi.nlm.nih.gov/genomes/all/GCA/000/001/405/GCA_000001405.15_GRCh38/seqs_for_alignment_pipelines.ucsc_ids/), and the resulting alignment records were sorted with Samtools (http://www.htslib.org/; version 1.7). Marking of PCR duplicates and calculations of insert-size and coverage information was performed using Picard (http://picard.sourceforge.net; version 2.17.10).

### Illumina mate-pair WGS

#### Library Preparation

Mate Pair libraries at insert sizes 2 kb, 5 kb, and 12 kb were generated from Venter/HuRef DNA using the Nextera Mate Pair Sample Preparation Kit (Illumina, Cat# FC-132–1001) following standard manufacturer’s instructions with the exception of the shearing step (see below). The Venter/HuRef DNA sample was first verified as high molecular weight (>15 kb) by running 60 ng, quantified by using the Qubit dsDNA HS Assay Kit (Life Technologies, Cat# Q32851), on 0.8% 1X TAE agarose gel next to the 1 kb Plus DNA Ladder (ThermoFisher Cat# 10787018). Afterwards, for each insert size, 4□μg of the high molecular weight genomic DNA was tagmented with biotinylated junction adapters and fragmented to about 7-8□kb on average in a 400 μL tagmentation reaction containing 12□μL of Tagmentase at 55□°C for 30□min. The tagmented DNA fragments were purified by adding 2X the volume of DNA Binding Buffer with Zymo Genomic DNA Clean & Concentrator Kit (Zymo Research, Cat# D4010) and eluted in 30 uL of Elution Buffer after two washes with the provided Wash Buffer. To fill in the gaps in the DNA adjacent to the junction adapters as a by-product of Tagmentation, single-strand displacement reaction was performed in a 200 μL reaction by adding 132 μL of water, 20 μL of 10x Strand Displacement Buffer, 8 μL of dNTPs, and 10 μL of Strand Displacement Polymerase to the 30 μL elution and at 20□°C for 30□min. DNA purification was then performed in 30 μL elution with 0.5x volume of AMPure XP Beads (Beckman Coulter, Cat# A63880) and size-selected by using BluePippin (Sage Science). The 0.75% DF 3-10kb Marker S1 – Improved Recovery and the 0.75% DF 10-18kb Marker U1 protocols were used for size selection on the BluePippin for insert sizes 5 kb and 12 kb respectively, and 0.75% DF 1-6kb Marker S1 protocol was used for insert size 2 kb. The “Tight Selection” option was used instead of “Range” for all size selections. The size selected DNA was then circularized overnight (12-16 hours) at 30□°C with Circularization Ligase in a 300□μL reaction.

After overnight circularization, the uncirculated linear DNA was digested by adding 9 μL of Exonuclease and incubated at 30□°C for 30 minutes and heat inactivated at 70□°C for 30 minutes. Afterwards, 12 μL of Stop Ligation Buffer was added. Circularized DNA was then transferred to T6 (6×32□mm) glass tube (Covaris, Part# 520031 and 520042) and sheared *twice* on the Covaris S2 machine (Intensity of 8, Duty Cycle of 20%, Cycles Per Burst of 200, Time of 40□s, Temperature of 2–6□°C). We find that shearing *twice* often creates a tighter final library size distribution which leads to a higher fraction of pass-filter clusters during the Illumina sequencing step. The mate pair fragments within the sheared DNA fragments contain the biotinylated junction adapter and were selected by binding to Dynabeads M-280 Streptavidin Magnetic Beads (Invitrogen, Part# 112-05D) by adding an equal volume of the Bead Bind Buffer (incubated at 20□°C for 15 minutes on shaking heat block at highest rpm setting). The non-biotinylated molecules in solution were washed away using the Wash Buffer. All downstream reactions were carried out on streptavidin beads with magnetic immobilization and washes with the Wash Buffer between successive reactions (e.g. End Repair, A-Tailing, and Adapter Ligation. The sheared DNA was first End-repaired followed by A-Tailing and TruSeq indexed adapter ligation. The adapter-ligated DNA was resuspended in 20 μL of Resuspension Buffer and then PCR amplified in a 50 μL reaction with 25 μL of PCR 2X Master Mix and 5 μL of Primers both provided in the Nextera Mate Pair Sample Preparation Kit (Illumina, Cat# FC-132–1001) to generate the final library. The thermocycling conditions are 98□°C/1□mm, 10 cycles of (98□°C/10□s, 60□°C/30□s, 72□°C/30□s), 72□°C/5□min, and 4□°C /hold. The 5 kb mate-pair library was PCR-amplified for 5 cycles instead of 10 cycles. For the 12 kb mate-pair library, 8 μg of input DNA was used instead of 4 ug. The amplified library (supernatant) was purified using a 0.66x volume of AMPure XP Beads (0.67x vol) and eluted in 20 μL of Resuspension Buffer. The size distribution of the library was determined by Agilent Technologies 2100 Bioanalyzer (High Sensitivity Assay), and the indexed library concentration was measured by the Qubit dsDNA HS Assay Kit (Life Technologies, Cat# Q32851).

#### Sequencing

The Mate-Pair libraries were sequenced on the Illumina NextSeq 500 using the NextSeq 500/550 Mid Output v2 kit (300 cycles) (Illumina, Cat# FC-404-2003) to generate 2×151□bp paired-end reads. The libraries were loaded onto the flowcell at a final concentration of 1.8pM and 1% PhiX Control v3 (Illumina, Cat# FC-110-3001). Additional rounds of sequencing also used a final library concentration of 1.8pM and 1% PhiX Control v3.

#### Analysis

Illumina Nextera Mate Pair junction adapter sequences were first trimmed using NxTrim ^22^ (version 0.4.3) with the “--aggressive --preserve-mp” settings in order to maximize the number of long-insert pairs. Nxtrim outputs four sets of reads, designated “Mate Pair”, “Paired-End”, “Singleton”, and “Unknown.” “Mate Pair” reads have junction adapter sequence trimmed off from the 3’ end of Read 1 and/or Read 2; “Paired-End” (short-insert) reads have junction adapter sequence trimmed from the 5’ end of Read 1 and/or Read 2; “Singleton” reads have junction adapter sequence trimmed from the middle of either Read 1 or Read 2 rendering one of the reads useless. “Unknown” reads have no junction adapter sequences detected. This is most likely because the junction adapter sequence sits in the un-sequenced portion of the template, thus whether reads are “Mate Pair” or “Paired-End” cannot be discerned. Nonetheless, mate-pair reads are present in the “Unknown” fractions as well as paired-end reads. The “Unknown” reads can be used for alignment and analysis if more long-insert information is desired ^22^. Here, the reads designated as “Mate Pair” and “Unknown” were combined, aligned with BWA-MEM ^23^ against the hg38 reference without ALT alleles (ftp://ftp.ncbi.nlm.nih.gov/genomes/all/GCA/000/001/405/GCA_000001405.15_GRCh38/seqs_for_alignment_pipelines.ucsc_ids/), and sorted using samtools (http://www.htslib.org/; version 1.7). Marking of PCR duplicates and calculations of insert-size and coverage information was performed using Picard (http://picard.sourceforge.net; version 2.17.10).

### 10X Genomics Chromium library for Illumina sequencing

#### Input genomic DNA preparation

The Venter/HuRef DNA sample (obtained from the Coriell Institute for Medical Research) was first verified as high molecular weight (>15 kb) by running 60 ng, quantified by using the Qubit dsDNA HS Assay Kit (Life Technologies, Cat# Q32851), on 0.8% 1X TAE agarose gel next to the 1 kb Plus DNA Ladder (ThermoFisher Cat# 10787018). Afterwards, 4□μg of the high molecular weight genomic DNA was loaded on a BluePippin (Sage Science) instrument to select for DNA fragments 30 kb to 80 kb using the “0.75%DF Marker U1 high-pass 30- 40 kb vs3” protocol. The concentration of the selected DNA fragments was then quantified by using the Qubit dsDNA HS Assay Kit (Life Technologies, Cat# Q32851) and diluted to 1 ng/ μL. The final dilution concentration of 1 ng/ μL was verified again by performing three technical replicates of Qubit dsDNA HS Assay with 5 μL of the DNA dilution as input. *Chromium whole-genome linked-read library preparation and sequencing* The linked-read whole-genome library was prepared using the Chromium Genome kit and reagent delivery system (10X Genomics, Pleasanton, CA). The linked-read library was made following standard manufacturer’s protocol with 10 cycles of PCR amplification. Briefly 1□ng of DNA (~300 genome equivalents) of size-selected high molecular DNA was partitioned into ~1.5 million oil droplets in emulsion, tagged with a unique 16 bp barcode within each droplet, and subjected isothermal amplification (30□°C for 3 hours; 65□°C for 30□minutes) by random priming within each droplet. Amplified (isothermal) DNA was then purified from the droplet emulsion following the manufacturer’s protocol using SPRI beads. The purified DNA was then End-Repaired and A-tailed followed by adapter ligation of adapter in the same reaction mixture. DNA was purified from the was the reaction mixture using SPRI beads and eluted in 40 uL. Sample Index PCR amplification (primers and 2X master mix provided in the Chromium Genome kit) was then performed on the eluted DNA in a toal volume of 100uL with the following thermocycling conditions: 98□°C/45□s, 10 cycles of (98□°C/20□s, 54□°C/30□s, 72□°C/20□s), 72□°C/1□min, and 4□°C/hold. Primer index SI-GA-A6 was used. DNA (final linked-read library) was purified from the PCR reaction with SPRI bead size selection following manufacturer’s protocol.

#### Sequencing

The final purified library was quantified by qPCR (KAPA Library Quantification Kit for Illumina platforms, Kapa Biosystems, Wilmington, MA) using the following thermocycling conditions: 95□°C/3 min, 30 cycles of (95□°C/5□s, 67□°C/30□s). The library concentration was calculated in nanomolar (nM) concentration and then diluted to 5 nM. Sequencing (2×151bp, 8 cycles of single indexing) on two lanes of Illumina HiSeq X (flowcell ID: H3MHGALXX, lanes #4 and #5) was performed at Macrogen (Rockville, MD) resulting in a total of 789,239,544 paired reads (**Table 1**).

#### Analysis

FASTQ files were generated from raw BCL files using “*mkfastq*” mode in the Long Ranger software (version 2.1.3) from 10X Genomics (Pleasanton, CA). 10X Genomics Chromium library index “SI-GA-A6” was specified in the required sample sheet file for “*mkfastq*”. Before alignment, the hg38 genome files were downloaded from ftp://ftp.ncbi.nlm.nih.gov/genomes/all/GCA/000/001/405/GCA_000001405.15_GRCh38/seqs_for_alignment_pipelines.ucsc_ids/GCA_000001405.15_GRCh38_no_alt_analysis_set.fna.gz and indexed using the “*mkref*” mode in Long Ranger. Sequencing alignment and haplotype phasing were performed using the “*wgs*” mode in Long Ranger, and the options *“--sex=male*” and *“--vcmode=freebayes*” were specified. Only “PASS” SNPs and Indels 50 bp or smaller were included in the final phased variant vcf.

## DATA RECORDS

The Venter/HuRef genome sequenced is publicly available through The Coriell Institute for Medical Research (Camden, NJ, USA) both as genomic DNA (catalog ID: NS12911) extracted from lymphoblastoid cell line (LCL) or as retroviral reprogrammed induced pluriplotent stem cell culture (catalog ID: GM25430). As described in the Methods, Venter/HuRef LCL DNA (NCBI SRA biosample accession SAMN03491120) was used for sequencing library preparation in this work.

### Illumina short-insert WGS

Approximately 100x sequencing coverage 2×100bp Illumina short-insert (200 bp) WGS data generated from the Illumina HiSeq 2000 is available through NCBI SRA accession SRR7097858 [Data Citation 1, **Table 5**]. Approximately 40x sequencing coverage 2×100bp Illumina short-insert (350 bp) WGS data generated from the Illumina HiSeq 2000 platform is available through NCBI SRA accession SRR7097859 [Data Citation 1, **Table 5**].

### Illumina mate-pair WGS

Illumina mate-pair data sequenced (2×150 bp) on the Illumina NextSeq 500 are available through NCBI SRA accessions SRR6951312, SRR6951313, and SRR6951310 for insert sizes 2 kb, 5 kb, and 12 kb respectively [Data Citation 1, **Table 5**].

### 10X Genomics Chromium linked-read Library

10X Genomics Chromium linked-read data sequenced (2×150 bp) on two lanes of the Illumina HiSeq X Ten is available through NCBI SRA accession SRR6951311 [Data Citation 1, **Table 5**]. The phased variants of the Venter/HuRef genome obtained through the analysis linked reads is available through dbSNP NCBI_ss# 2137543904 to 3651364986 (For phasing information, request for original submitted vcf file through NCBI dbSNP.) [Data Citation 2: NCBI dbSNP NCBI_ss# 2137543904-3651364986].

## TECHNICAL VALIDATION

### Illumina short-insert WGS

Sequencing quality of the WGS mate-pair libraries were assessed using FastQC (**Supplementary Information**). Insert-size, coverage, GC-bias, alignment, and duplication metrics were analyzed using Picard tools (http://broadinstitute.github.io/picard/). These statistics are summarized in Table 1, Table 2 and Figure 2A.

### Illumina mate-pair WGS

Sequencing quality of the WGS mate-pair libraries was assessed using FastQC (**Supplementary Information**). Insert-size, coverage, GC-bias, alignment, and duplication metrics were analyzed using Picard tools (http://broadinstitute.github.io/picard/). These statistics are summarized in **Table 1, Table 2 and Figure 2c-j**. Read fractions that were designated by NxTrim ^22^ as “Mate Pair”, “Paired-End”, “Singletons”, and “Unknown” are summarized in **Table 3**. The “Mate Pair” fraction for all libraries fall within the expected range (~40-60%). Expected for mate-pair libraries, the relatively high rates of PCR duplication (**Supplementary Information**) result in significant decreases in sequence coverage (3x to 7x) (**Table 1, Table 2, Figure 2**). However, the more useful metric for mate-pair sequencing is high physical coverage ^24^. Here, physical coverage (CF) is calculated by the equation C = C^R^ x C^F^ where C is the sequencing coverage and C^R^ is the mean fractional coverage of input DNA fragments. The mean insert sizes for the mate pair libraries are 1.8 kb, 4.8 kb, and 12.2 kb (**Table 2, Figure 2**), which results in physical coverage values of 49x, 122x, and 145x respectively. For the 2 kb mate-pair library for an example, C is 7x, and C^R^ is 0.14 or (130 bp + 131 bp)/1845 bp, thus CF is 49x (**Table 2**). The average final library fragment lengths were approximately 800 bp, 800 bp, and 500 bp for the 2 kb, 5 kb and 12 kb mate-pair libraries respectively. The differences in average library fragment lengths most likely contributed to the more extreme tails of the normalized coverage vs GC% for the 2 kb and 5 kb mate-pair libraries (Figure 2e, g) ^25^.

### 10X Genomics Chromium Library

Sequencing quality of the linked-read library was assessed using FastQC (**Supplementary Information**). Input molecule length, coverage, alignment, duplication, droplet barcode, and phasing metrics were analyzed using the Long Ranger software version 2.1.5 ^26^ (**Table 1, Table 2, Table 4 and Figure 3)**. Overall, 2.4 million and 1.5 million, 0.42 million and 0.29 million heterozygous and homozygous SNVs and indels respectively were called (**Table 4**). Of which, 96.7% and 93.85% of heterozygous SNVs and Indels respectively were successfully phased in the Venter/HuRef Genome in a total of 8882 haplotype blocks (N50 ~ 0.9 Mbp, longest phase block ~ 6.5 Mbp) (**Table 4**). Phase blocks for each chromosome are shown in **Figure 4**. Similar to mate-pair libraries, the physical coverage of the linked read library is calculated to be 128x from the mean input DNA molecule length of 32kb.

**Figure 3.**
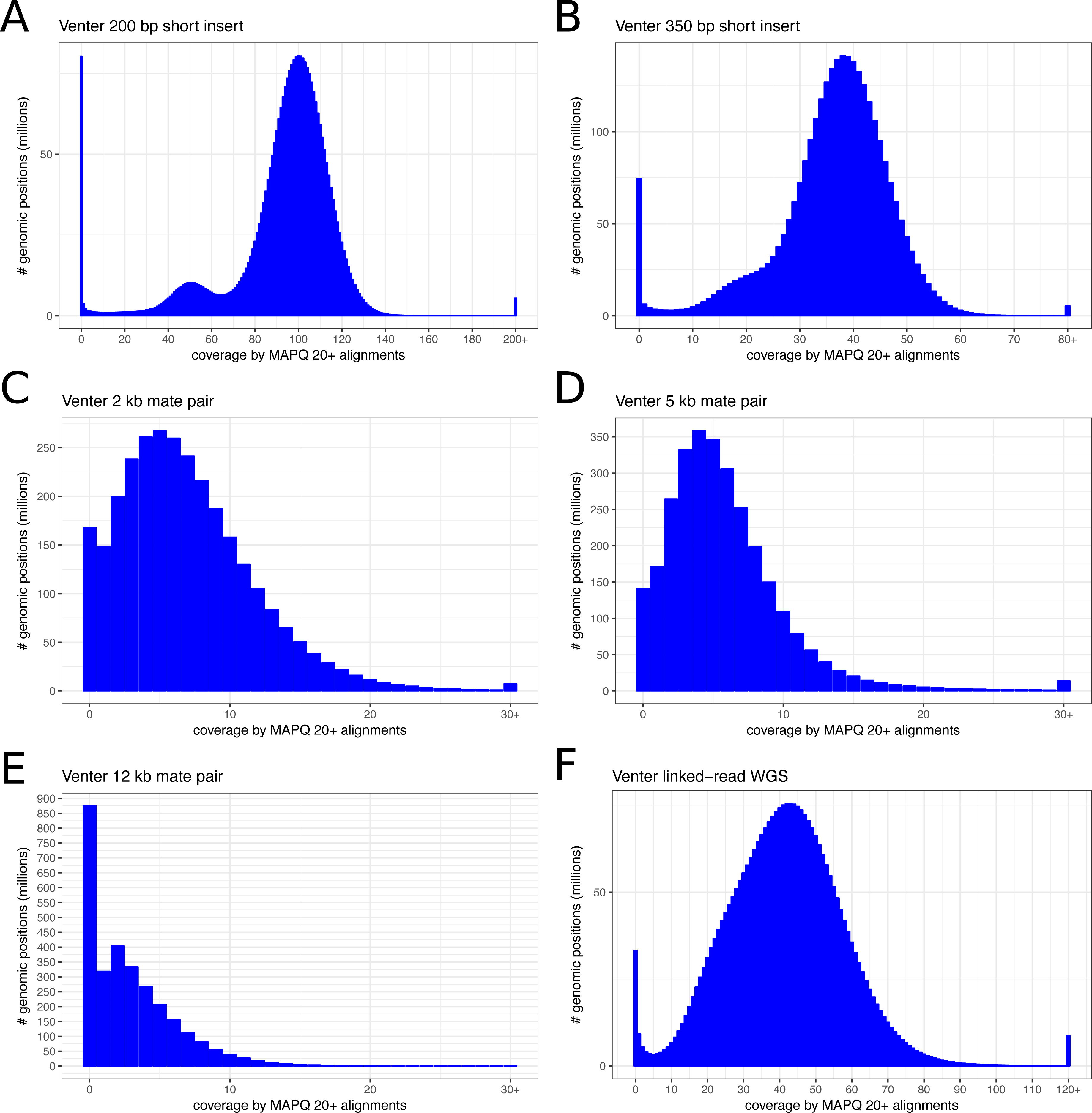
Coverage (deduplicated) histograms. **(a, b)** short-insert, **(c, d, e)** 2 kb, 5 kb, and 12 kb mate-pair, and **(f)** linked-read libraries. Only reads with mapping score > 20 were used.

**Figure 4.**
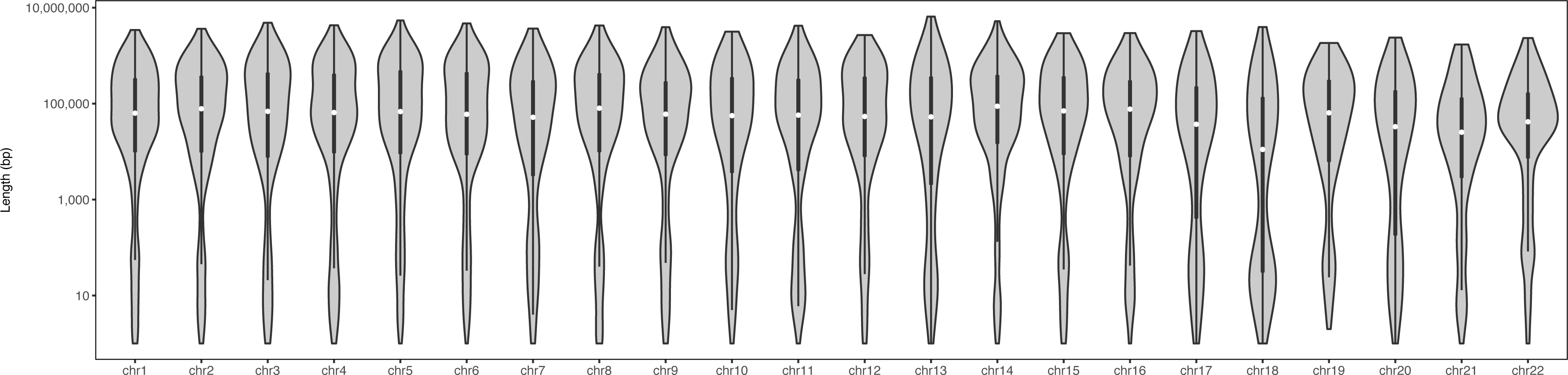
Violin plot of sizes of haplotype blocks constructed using linked-read sequencing (128x physical coverage) for HuRef/Venter Genome for all chromosomes.

## USAGE NOTES

The Venter/HuRef genome sequenced in this work is publicly available as both cell line and DNA from Coriell Institute for Medical Research. The mate-pair and linked-read sequencing data used the same DNA sample/extraction as input. It is possible that small differences may exist when compared to the short-insert datasets since the input DNA came from different cell passages and extractions. Researchers are especially encouraged to use the sequencing data in this work in combination with diploid Sanger sequencing data available for the Venter/HuRef genome published in Levy et al ^17^.

## ACKNOWLEDGEMENTS

A.E.U. was supported by NIH grant P50-HG007735 and the Stanford Medicine Faculty Innovation Program, and B.Z. was additionally supported by NIH training grant T32- HL110952-04. W.H.W. was supported by NIH grants P50-HG007735 and R01- HG007834. J.G.A. received funding from NIH training grant T32-GM096982 and NSF Graduate Fellowship DGE-114747.

## AUTHOR CONTRIBUTIONS

BZ and RP performed the experiments. JGA, BZ, SSH, and YH performed the data analysis. BZ, JGA, WHW and AEU conceived of the work and wrote the manuscript.

---

**Table 1. Summary of library construction and sequencing for short-insert, mate-pair, and linked-read HuRef/Venter WGS libraries.**

**Table 2. Summary of post sequencing QC, alignment, duplication, coverage and insert-size analysis for all libraries.**

**Table 3. Statistics for trimming of Nextera junction adapter sequence using NxTrim for all mate-pair libraries.**

**Table 4. Summary of metrics for linked-read sequencing and phasing of the HuRef/Venter genome.**

**Table 5. Details of Data Citation 1 (SRP137779).** * obtained from Mu et al ^20^. Phred-33 encoding for all files.

^*^For phasing information, request the originally submitted VCF file through NCBI dbSNP.

